# KSR1 knockout mouse model demonstrates MAPK pathway’s key role in cisplatin- and noise-induced hearing loss

**DOI:** 10.1101/2023.11.08.566316

**Authors:** Matthew A. Ingersoll, Richard D. Lutze, Regina G. Kelmann, Daniel F. Kresock, Jordan D. Marsh, Rene V. Quevedo, Jian Zuo, Tal Teitz

## Abstract

Hearing loss is a major disability in everyday life and therapeutic interventions to protect hearing would benefit a large portion of the world population. Here we found that mice devoid of the protein kinase suppressor of RAS 1 (KSR1) in their tissues (germline KO mice) exhibit resistance to both cisplatin- and noise-induced permanent hearing loss compared to their wild-type KSR1 littermates. KSR1 is expressed in the cochlea and is a scaffold protein that brings in proximity the mitogen-activated protein kinase (MAPK) proteins BRAF, MEK and ERK and assists in their activation through a phosphorylation cascade induced by both cisplatin and noise insults in the cochlear cells. Deleting the KSR1 protein tempered down the MAPK phosphorylation cascade in the cochlear cells following both cisplatin and noise insults and conferred hearing protection of up to 30 dB SPL in three tested frequencies in mice. Treatment with dabrafenib, an FDA-approved oral BRAF inhibitor, downregulated the MAPK kinase cascade and protected the KSR1 wild-type mice from both cisplatin- and noise-induced hearing loss. Dabrafenib treatment did not enhance the protection of KO KSR1 mice, as excepted, providing evidence dabrafenib works primarily through the MAPK pathway. Thus, either elimination of the KSR1 gene expression or drug inhibition of the MAPK cellular pathway in mice resulted in profound protection from both cisplatin- and noise-induce hearing loss. Inhibition of the MAPK pathway, a cellular pathway that responds to damage in the cochlear cells, can prove a valuable strategy to protect and treat hearing loss.

**Significance Statement:** Ten percent of the world population suffers from hearing loss but this impairment may be preventable. We show that mice devoid of the KSR1 protein (KO) exhibit resistance to cisplatin- and noise-induced permanent hearing loss compared to wild-type littermates that harbor the protein. Removing KSR1 tempers down the MAPK phosphorylation cascade of BRAF-MEK-ERK induced in the cochlea following cisplatin and noise insults. Treatment of KSR1 wild-type mice following cisplatin or noise with an FDA-approved BRAF inhibitor, dabrafenib, protected the hearing and, importantly, did not confer additional protection to the KO KSR1 mice. Hence, the MAPK pathway has a unique role in responding to cochlear damage, and removing KSR1 gene expression or drug inhibition of the pathway results in hearing protection.

## Introduction

Irreversible hearing loss afflicts more than 10% of the world population; however, there is only one Food and Drug Administration (FDA) approved drug to prevent any type of hearing loss^1,2^. Sodium Thiosulfate (brand name ®PEDMARK) was recently approved by the FDA for the prevention of cisplatin-induced hearing loss in pediatric patients with non-metastatic tumors^3–5^. No other patient population or type of hearing loss has a drug that can protect from this very common disability; therefore, there is a dire need to find compounds that protect from different types of hearing loss. Our laboratory has shown that inhibitors of the mitogen activated protein kinase (MAPK) pathway protect mice from cisplatin and noise-induced hearing loss^6-,8^. Other laboratories have also demonstrated that activation of the MAPK pathway occurs after ototoxic insult and targeting this pathway can be a therapeutic approach to protect from hearing loss^9–15^. Dabrafenib, a BRAF inhibitor, is a promising otoprotective drug in preclinical studies which has been studied by our laboratory^6,8^. Dabrafenib, which is well tolerated by patients, is already FDA approved for cancer treatment which makes it a promising compound to repurpose for protection from cisplatin and noise-induced hearing loss^16,17^. Dabrafenib has demonstrated significant protection from cisplatin ototoxicity in three mouse strains and multiple cisplatin treatment regimens^6,8^. Additionally, it protects mice from noise-induced hearing loss which indicates it is not limited as an otoprotective agent to one type of damaging stimulus^6^. Dabrafenib is a specific BRAF inhibitor, but the mechanistic target of its protective effect in hearing loss has not yet been confirmed.

The MAPK pathway is involved in a multitude of critical cellular processes with RAF, MEK, and ERK as three of the main kinases in the pathway^18^. Kinase Suppressor of Ras 1 (KSR1) is a scaffolding protein that brings together RAF, MEK, and ERK to be rapidly phosphorylated^19^. Kinases in the MAPK pathway are activated when phosphorylated and KSR1 KO mice have been shown to have reduced MAPK activation^19–21^. It has been demonstrated that KSR1 is needed for optimal MAPK signaling to occur^19^. The MAPK pathway has typically been associated with cell proliferation and survival^22^; however, activation of the pathway has also been shown to lead to cellular stress and eventually cell death^23^. The type of stimulus, the cell type, and the duration of MAPK activation all play a role in determining whether it leads to cell survival or cell stress^6,23–25^. MAPK signaling has been implicated in cisplatin and noise-induced hearing loss^6–8,10,11^, and here we show that the KSR1 KO mice do not have cochlear structural abnormalities and exhibit normal hearing before damage; therefore, utilization of the KSR1 KO mouse model can help elucidate the role of the MAPK pathway in noise and cisplatin-induced hearing loss while germline knockouts of RAF, MEK, and ERK are embryonically lethal^26^. The KSR1 KO genetic mouse model will also be a useful tool to determine whether dabrafenib is exhibiting its otoprotective effect through the MAPK pathway or some other off target effects.

In this study, we validate that activation of the MAPK pathway leads to both cisplatin and noise-induced hearing loss by utilizing the KSR1 KO mouse model. We show that MAPK genes are ubiquitously expressed throughout many of the cell types in the inner ear and KSR1 KO mice had decreased MAPK signaling compared to WT littermates. KSR1 KO mice were resistant to cisplatin and noise-induced hearing loss compared to WT littermates while also being resistant to the increase in ERK activation following ototoxic insult that was demonstrated in WT mice. Additionally, dabrafenib was orally administered to KO and WT littermates to confirm that its protective mechanism of action was through MAPK inhibition. This study shows that stress-induced activation of the MAPK pathway in the cochlea causes permanent damage and that dabrafenib protects from hearing loss through inhibition of the MAPK pathway.

## Results

### Expression of MAPK genes in C57BL/6 mouse cochlea

To gain insight on the expression of MAPK genes throughout the cochlea, we analyzed Xu et al’s scRNA dataset from postnatal day 28 (P28) C57BL/6 mice, the same strain of mice used to generate the genetic KSR1 knockout model^27^. Based on genetic profiling, 6,386 cells from two biological replicates were organized into cell-type clusters by differentially expressed genes (DEGs) and visualized by UMAP plot (Figure 1A). The dataset was then examined for MAPK gene expression to generate gene-specific UMAP plots and expression-pattern violin plots for the following: Braf (BRAF), Map2k1 (MEK1), Map2k2 (MEK2), Mapk3 (ERK1), Mapk1 (ERK2), Ksr1 (KSR1), and Ksr2 (KSR2) (Figure 1B-C). Braf and Map2k1 have a similar ubiquitous expression pattern throughout the cochlea at moderate levels, with notable densities in OHCs, basal cells, inner/outer sulcus cells, and inner boarder cells. Map2k2, Mapk3, and Mapk1 are also ubiquitously expressed throughout the cochlea, though at higher levels, with notable densities in OHCs, basal cells, intermediate cells, Schwann cells, root/spindle cells, inner phalangeal cells, inner/outer sulcus cells, pillar/dieter cells, and inner boarder cells. Furthermore, Ksr1 was notably expressed only in specific cell types including OHCs, inner border cells, inner phalangeal cells, fibrocytes, intermediate cells, inner/outer sulcus cells, root/spindle cells, and Reisner’s membrane. Interestingly, Ksr2 had minimal expression throughout the cochlea with a moderate density only observed in OHCs.

**Figure 1:**
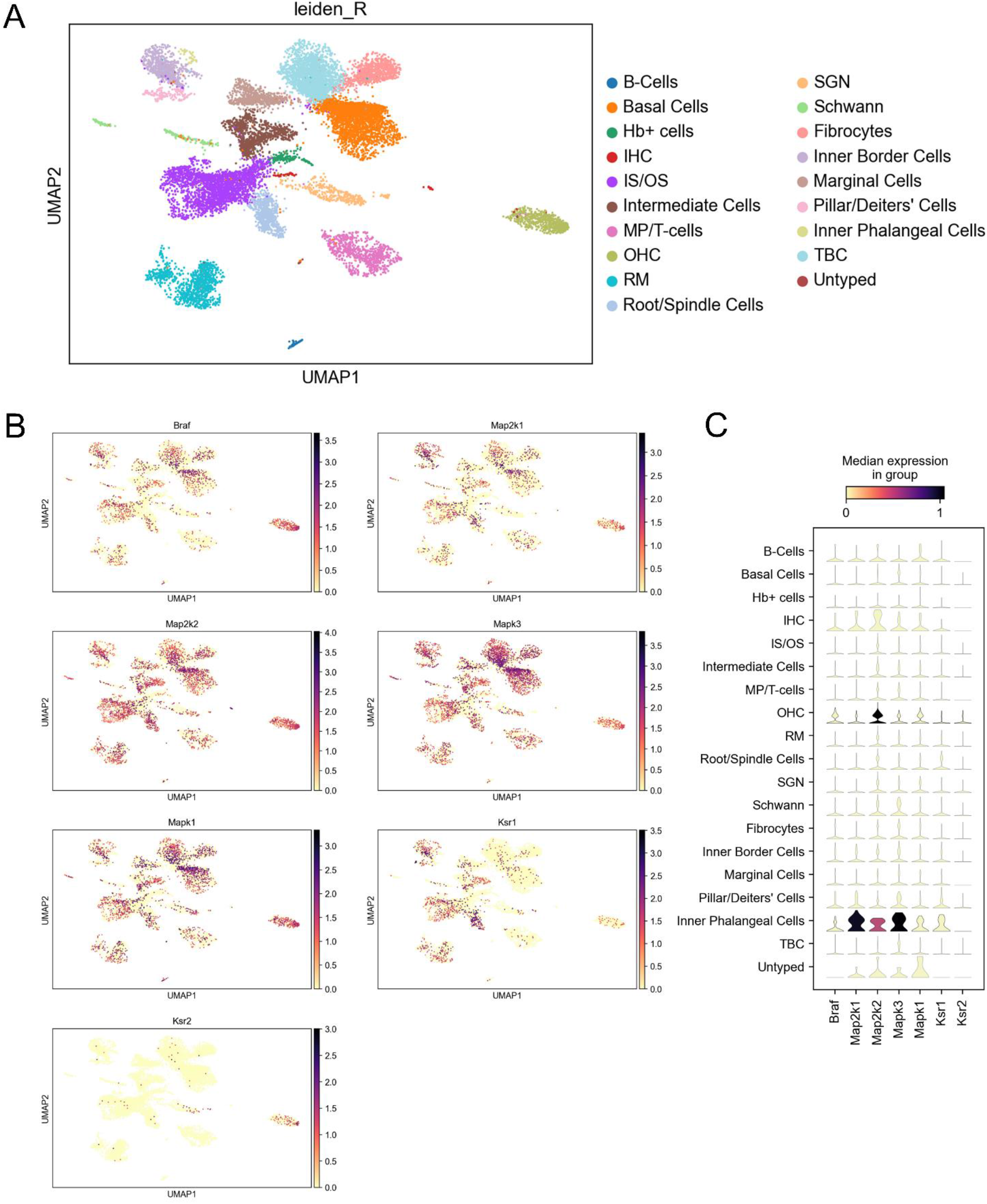
Single Cell RNA-Sequence expression of MAPK proteins in adult P28 C57BL/6 mouse cochlea. (**A**) UMAP plot of ∼18,000 cochlear cells from P28 mice organized into 19 cell-type clusters. Abbreviations: Inner Hair Cells (IHCs), Outer Hair Cells (OHCs), Hemoglobin-positive cells (HB+), Inner Sulcus/ Outer Sulcus (IS/OS), Reissner’s Membrane (RM), Spiral Ganglion Neurons (SGNs), Tympanic Boarder Cells (TBCs). (**B**) Feature plots showing the expression of Braf, Map2k1 (MEK1), Map2k2 (MEK2), Mapk3 (ERK1), Mapk1 (ERK2), Ksr1, and Ksr2 genes for the same cells in (A). (**C**) Violin plots showing median gene expression from (B) across cell-type clusters.

### Characterization of the KSR1 genetic KO model

Homozygous KSR1 WT/WT (WT) and KO/KO (KO) mice were characterized to examine differences in MAPK protein phosphorylation and activation, as well as determine whether there were any differences in cochlear morphology or hearing ability between the two genotypes. Due to the low level of KSR1 protein in the mouse cochlea, the combined organ of Corti lysates of 3 WT or 3 KO mice were pooled and analyzed by western blot to confirm that the protein is expressed in WT mice, but not KO mice (Figure 2A). Next, immunoblot was performed for individual organ of Corti lysates from 3 WT or 3 KO mice for total and phosphorylated/activated protein levels of BRAF, MEK1/2, and ERK1/2 and blot intensities of phosphorylated protein relative to loading control (α-tubulin) were quantified (Figure 2B-C). Levels of total and phosphorylated BRAF, MEK1/2, and ERK1/2 protein were lower in KSR1 KO mice compared to WT mice. When quantified, levels of phosphorylated BRAF and MEK1/2 trended lower in KO mice, while levels of phosphorylated ERK1/2 were significantly lower in KO mice compared to WT mice. In addition, cross-sections of WT and KO mouse cochleae were stained with Hematoxylin and Eosin (H&E) and examined under a light microscope. No differences in structural morphology were found (Figure 2D). Finally, functional hearing tests, including auditory brainstem response (ABR) and distortion product otoacoustic emission (DPOAE), were performed on P42 WT and KO mice and no differences in baseline hearing ability were detected between the two genotypes (Figure 2E-F).

**Figure 2:**
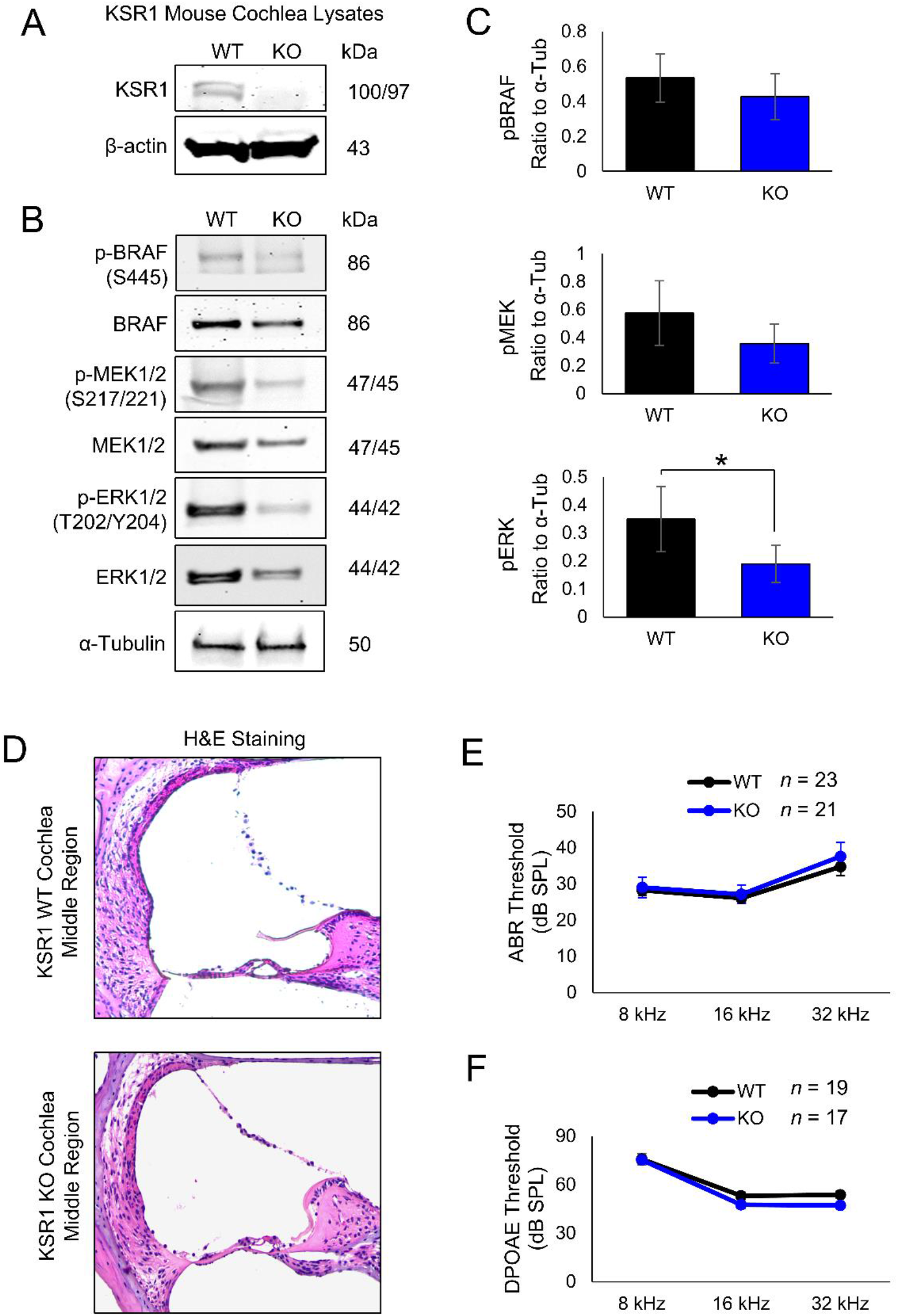
KSR1 KO C57BL/6 mice have reduced MAPK phosphorylation and normal hearing function. (**A**) KSR1 and β-actin loading control Western blot images of organ of Corti lysates from pooled lysates of 3 WT and 3 KO P42 mice. (**B**) Representative Western blot images of BRAF, MEK, and ERK phosphorylation from KSR1 WT and KO organ of Corti lysates, α-tubulin loading control, *n* = 3. (**C**) ImageJ band intensity quantification of (B), normalized to α-tubulin. Data shown as means ± SEM, \**P* < 0.05 by t-test. (**D**) Representative H&E staining of middle-turn cochlear region of KSR1 WT and KO mice, *n* = 3. (**E**) ABR thresholds of KSR1 WT (black) and KO (blue) P42 mice. (**F**) DPOAE thresholds of KSR1 WT (black) and KO (blue) P42 mice.

### KSR1 KO and dabrafenib treated mice are resistant to cisplatin-induced hearing loss

Previous studies indicate inhibition of the MAPK pathway by dabrafenib confers protection from cisplatin-induced hearing loss and we sought to determine whether genetic KO of KSR1 and suppression of the pathway provides similar protection^6,8^. Initially, baseline ABR thresholds were determined for P42 KSR1 WT and KO mice, mice were allowed to recover for 7 days and then administered a single, high-dose intraperitoneal injection of cisplatin of 18 mg/kg, an optimized dose for hearing loss in C57/BL/6 mice. Mice were then allowed to recover for 14 days before ABR was again performed (Figure 3A). KSR1 WT and KO raw pre- and post-cisplatin ABR thresholds are shown in Figure 3B while ABR threshold shifts are shown in Figure 3C. Cisplatin treatment induced significant ABR threshold shifts in KSR1 WT mice of 27, 25, and 35 decibels (dB) at 8, 16, and 32 kHz frequencies respectively when compared to KO mice with only 3, 4, and 8 dB hearing loss at 8, 16, and 32 kHz. Similarly, post-cisplatin ABR wave 1 amplitudes at 16 kHz were lower in WT mice compared to KO mice, with a significant difference observed at 90 and 70 dB (Figure 3D). After final hearing tests were concluded, mouse cochleae were collected and examined for OHC death, which cisplatin is known to cause. Representative images of myosin VI-stained whole mount cochleae are shown and quantification of OHCs per 160 µm reveal WT mice lost significantly more OHCs than KO mice in the middle and basal regions (Figure 3E-F). While WT mice had an average of 42 and 40 OHCs in middle and basal regions respectively, KO mice maintained an average of 64 and 61.

**Figure 3:**
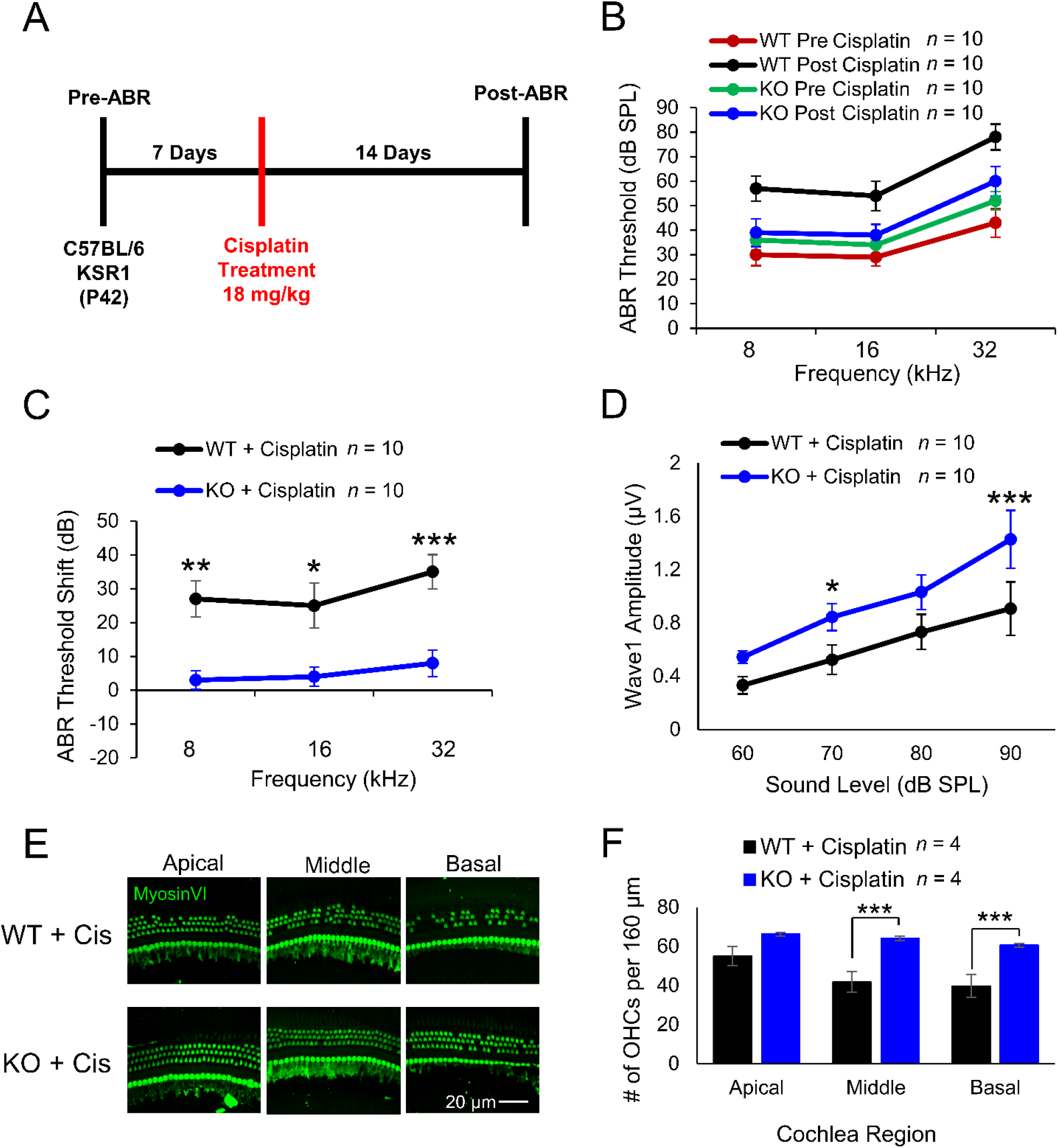
KSR1 KO mice are resistant to cisplatin-induced hearing loss. (**A**) Schedule of auditory testing and 18 mg/kg cisplatin IP administration for KSR1 WT and KO adult P42 mice. (**B**) ABR thresholds of mice before (WT-red, KO-green) and after (WT-black, KO-blue) cisplatin administration following protocol (A). (**C**) ABR threshold shifts for KSR1 WT (black) and KO (blue) mice calculated from (B). (**D**) Post-ABR wave 1 amplitudes from (B) at 16 kHz. (**E**) Representative confocal images of myosin VI stained KSR1 WT and KO apical, middle, and basal cochlear regions 14 days after cisplatin treatment. (**F**) Quantification of OHCs from (E) per 160 µm in apical, middle, and basal cochlear regions. Data shown as means ± SEM, \**P* < 0.05, \*\**P* < 0.01, \*\*\**P* < 0.001 by two-way ANOVA with Bonferroni post hoc test.

To determine whether dabrafenib confers protection from cisplatin-induced hearing loss through inhibition of the MAPK pathway, KSR1 WT and KO mice were again treated using the single high-dose cisplatin protocol with ABR readings performed before and after the experiment. Dabrafenib was administered by oral gavage at 12 mg/kg, twice daily for three days, with the first treatment given 45 mins prior to 18 mg/kg cisplatin injection (Figure 4A). The cohorts consisted of dabrafenib treated WT and KO mice, cisplatin treated WT and KO mice, and WT and KO mice cotreated with dabrafenib and cisplatin. Compared to all other cohorts, WT mice treated with cisplatin had significantly elevated average ABR threshold shifts of 27, 23, and 34 dB at 8, 16, and 32 kHz respectively. WT mice cotreated with dabrafenib and cisplatin, KO mice treated with cisplatin, and KO mice cotreated with dabrafenib and cisplatin had similar average ABR threshold shifts of 8, 8, and 5 dB at 8 kHz respectively, 6, 5, and 7 dB at 16 kHz, and 14, 11, and 9 dB at 32 kHz. Mice of both genotypes treated with dabrafenib alone experienced no change in ABR thresholds (Figure 4B). Additionally, post-experimental wave 1 amplitudes at 16 kHz followed a similar trend in which WT mice treated with cisplatin had significantly lower amplitudes than all other cisplatin treated cohorts at 90 dB as well as KO mice treated with cisplatin at 70 dB (Figure 4C). Finally, OHC counts were again performed on cochlea collected after the protocol concluded. KSR1 WT mice treated with cisplatin were found to possess significantly fewer OHCs than all other cisplatin treated cohorts at the middle and basal regions, as well as fewer OHCs than WT mice cotreated with cisplatin and dabrafenib and KO mice treated with cisplatin at the apical region (Figure 4D-E).

**Figure 4:**
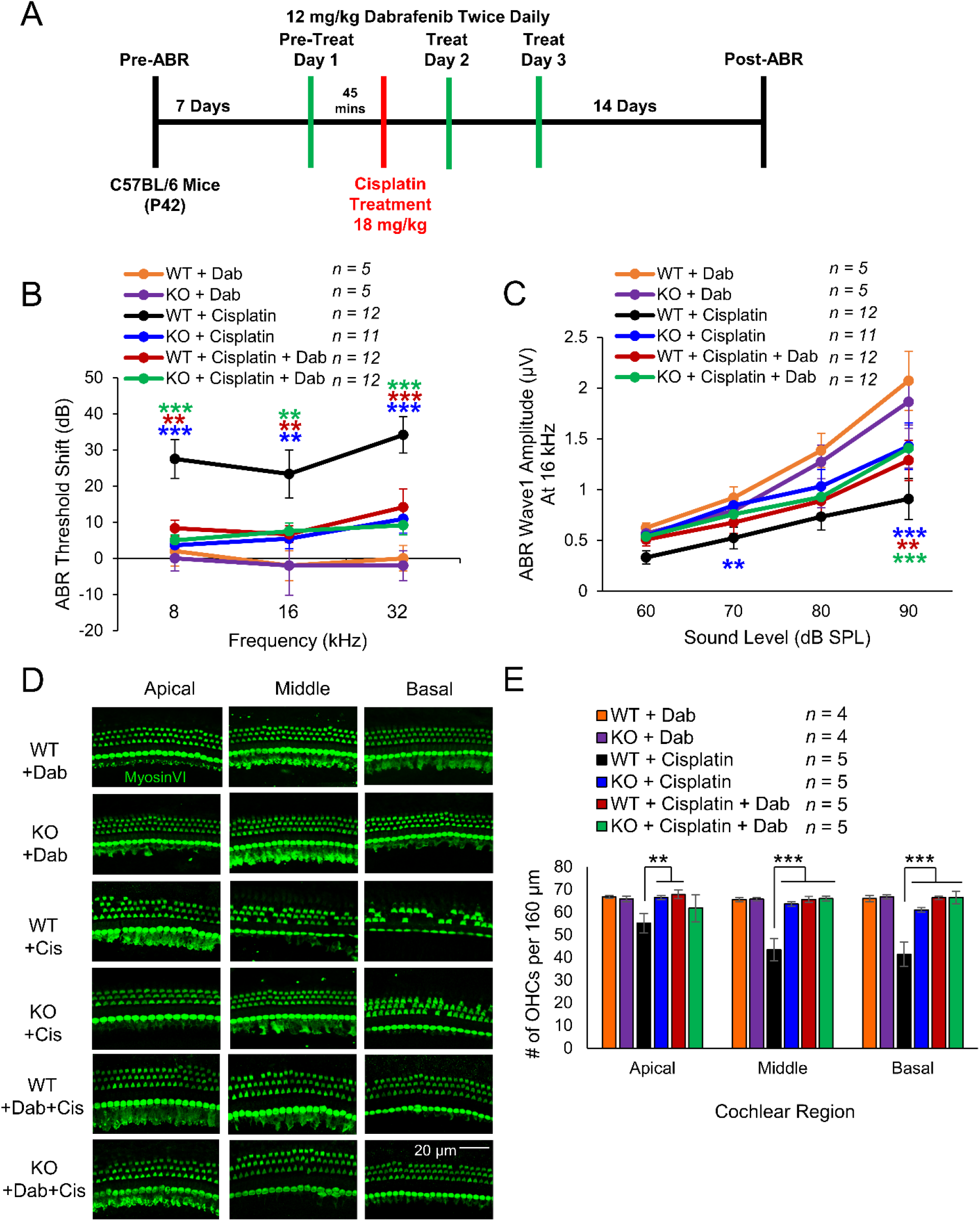
Dabrafenib confers protection from cisplatin-induced hearing loss to KSR1 WT, but not KO mice. (**A**) Schedule of auditory testing and administration of 12 mg/kg twice daily dabrafenib by oral gavage and 18 mg/kg cisplatin IP injection for KSR1 WT and KO adult P42 mice. (**B**) ABR threshold shifts following protocol in (A). WT (black) and KO (blue) mice treated with cisplatin alone, WT (orange) and KO (purple) mice treated with dabrafenib alone, WT (red) and KO (green) mice cotreated with dabrafenib and cisplatin. (**C**) Post- ABR wave 1 amplitudes from (B) at 16 kHz. (**D**) Representative confocal images of myosin VI-stained apical, middle, and basal cochlear regions from mice in (B). (**E**) Quantification of OHCs from (D) per 160 µm in apical, middle, and basal cochlear regions. Data shown as means ± SEM, \**P* < 0.05, \*\**P* < 0.01, \*\*\**P* < 0.001 by two- way ANOVA with Bonferroni post hoc test.

### KSR1 KO and dabrafenib treated mice are resistant to noise-induced hearing loss

Similar to dabrafenib’s protection against cisplatin, dabrafenib’s noise protection has previously been suggested to occur through inhibition of the MAPK pathway and the KSR1 KO model is here used to determine whether genetic suppression of the MAPK pathway offers similar protection^6^. Initially, baseline ABR and DPOAE thresholds were determined for P42 KSR1 WT and KO mice, mice were allowed to recover from anesthetic for 7 days and then subjected to 100 dB 8-16 kHz octave band noise for two hours. Mice were then allowed to recover for 14 days before ABR and DPOAE were again performed (Figure 5A). KSR1 WT and KO raw pre- and post-noise exposure ABR thresholds are shown in Figure 5B while ABR threshold shifts are shown in Figure 5C. Noise exposure induced significant average ABR threshold shifts in WT mice of 32, 43, and 34 decibels (dB) at 8, 16, and 32 kHz frequencies respectively when compared to KO mice with only 5, 12, and 8 dB threshold shifts at 8, 16, and 32 kHz. Post-noise exposure average wave 1 amplitudes at 16 kHz were then quantified and WT mice were found to have significantly lower amplitudes compared to KO mice at 90 dB (Figure 5D). Moreover, WT mice also had significantly larger DPOAE threshold shifts compared to KO mice with 25 and 8 dB respectively at 16 kHz as well as 15 and 0 dB at 32 kHz (Figure 5E). Due to the resilience against hearing loss of KSR1 KO mice when exposed to 100 dB of noise, the experiment was repeated, and mice were exposed to 106 dB noise exposure in the 8-16 kHz octave band for 2 hours. Again, WT mice experienced significantly greater hearing loss at all frequencies tested with threshold shifts of 41, 50, and 45 dB at 8, 16, and 32 kHz with respect to KO animals with threshold shifts of 23, 31, and 31 dB (Figure 5F). Finally, to examine differences in synaptopathy, cochleae from animals exposed to 100 dB of noise were stained with hair cell marker myosin VI and Ctbp2 antibody to identify pre-synaptic puncta. The number of Ctbp2 puncta per IHC in the 16 kHz region were quantified and WT mice were found to possess ∼10 puncta per IHC while KO mice maintained significantly more puncta with ∼14 per cell (Figure 5G-H).

**Figure 5:**
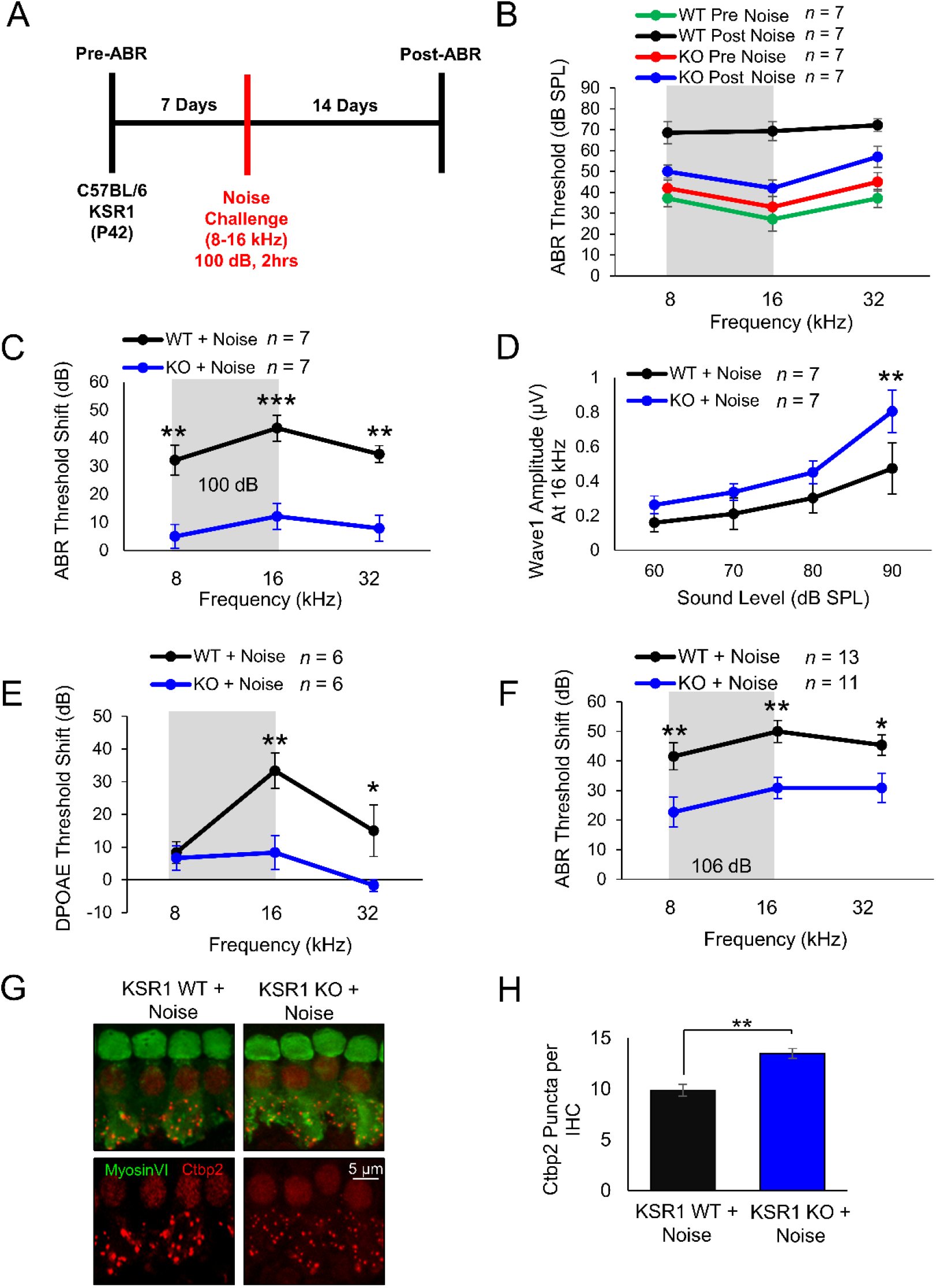
KSR1 KO mice are resistant to noise-induced hearing loss. (**A**) Schedule of auditory testing and noise exposure for KSR1 WT and KO adult P42 mice. (**B**) ABR thresholds of mice before (WT-red, KO-green) and after (WT-black, KO-blue) 2 hr 100 dB 8-16 kHz noise exposure following protocol (A). (**C**) ABR threshold shifts for KSR1 WT (black) and KO (blue) mice calculated from (B). (**D**) Post-ABR wave 1 amplitudes from (B) at 16 kHz. (**E**) DPOAE threshold shifts for KSR1 WT (black) and KO (blue) mice after 2 hr 100 dB 8-16 kHz noise exposure. (**F**) ABR threshold shifts of KSR1 WT (black) and KO (blue) mice after 2 hr 106 dB 8-16 kHz noise exposure following protocol (A). Data shown as means ± SEM, \**P* < 0.05, \*\**P* < 0.01, \*\*\**P* < 0.001 by two-way ANOVA with Bonferroni post hoc test. (**G**) Representative confocal images of Myosin VI (green) and Ctbp2 (red) middle-turn cochlear regions from mice in (B). (**H**) Quantification of average Ctbp2 puncta per IHC from (E) in middle cochlear region, 12 IHCs per cochlea, *n* = 5. Data shown as means ± SEM, \*\**P* < 0.01 by one-way ANOVA with Bonferroni post hoc test.

Next, the 100 dB noise protocol was repeated with the addition of dabrafenib treated cohorts to evaluate whether dabrafenib provided protection from noise induced hearing loss through inhibition of the MAPK pathway. Noise damage often cannot be predicted, so dabrafenib treatment began 24 hours after noise insult. The compound was administered by oral gavage at 12 mg/kg, twice daily, for three days (Figure 6A). The cohorts consisted of dabrafenib treated WT and KO mice, noise-exposed WT and KO mice, and noise-exposed WT and KO mice treated with dabrafenib. Compared to all other cohorts, WT mice exposed to noise had significantly elevated average ABR threshold shifts of 31, 44, and 37 dB at 8, 16, and 32 kHz, respectively. Noise-exposed WT mice treated with dabrafenib, noise-exposed KO mice, and noise-exposed KO mice treated with dabrafenib had similar average ABR threshold shifts of 9, 3, and 8 dB at 8 kHz respectively, 11, 14, and 9 dB at 16 kHz, and 6, 4, and 4 dB at 32 kHz. Mice of both genotypes treated with dabrafenib alone experienced no change in ABR thresholds (Figure 6B). In addition, WT mice exposed to noise had significantly lower wave 1 amplitudes at 16 kHz compared to all other noise-exposed cohorts at 90 and 80 dB (Figure 6C). To determine dabrafenib’s effect on synaptopathy, quantification of the average number of Ctbp2 puncta per IHC in the 16 kHz region was again performed. Noise-exposed WT mice had significantly reduced number of puncta with, ∼10 per cell, compared to all other cohorts. While dabrafenib offered significant protection to WT mice exposed to noise, the drug had no significant additional effect on noise-exposed KO mice (Figure 6D-E).

**Figure 6:**
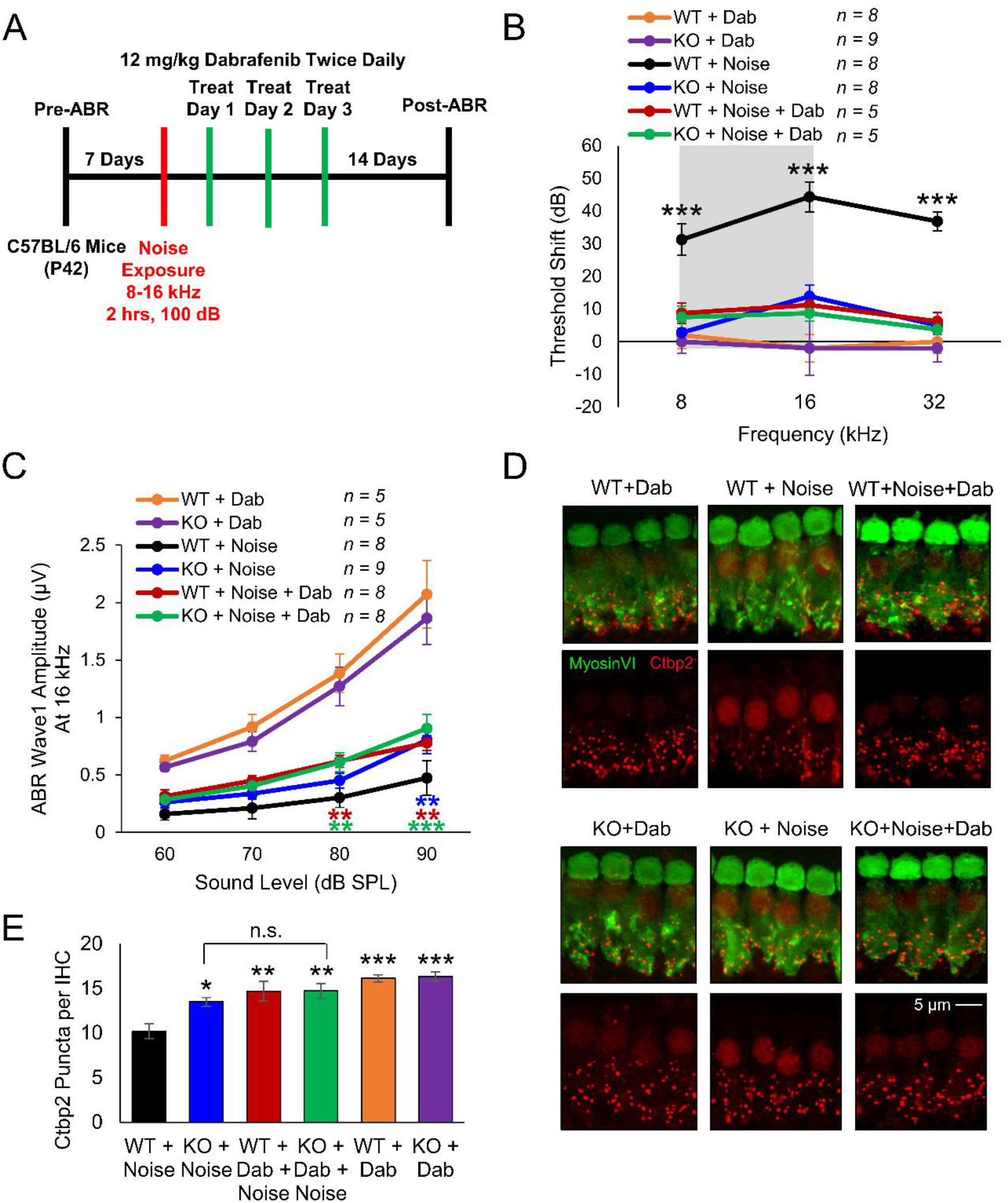
Dabrafenib confers protection from noise-induced hearing loss to KSR1 WT, but not KO mice. (**A**) Schedule of auditory testing, 2 hr 100 dB 8-16 kHz noise exposure, and 12 mg/kg dabrafenib administration for KSR1 WT and KO adult P42 mice. (**B**) ABR threshold shifts following protocol in (A). WT (black) and KO (blue) mice exposed to noise, WT (orange) and KO (purple) mice treated with dabrafenib alone, WT (red) and KO (green) mice treated with dabrafenib and exposed to noise. (**C**) Post-ABR wave 1 amplitudes from (B) at 16 kHz. Data shown as means ± SEM, \*\**P* < 0.01, \*\*\**P* < 0.001 by two-way ANOVA with Bonferroni post hoc test. (**D**) Representative confocal images of Myosin VI (green) and Ctbp2 (red) stained middle-turn cochlear regions from mice in (B). (**E**) Quantification of average Ctbp2 puncta per IHC from (D) in middle cochlear region, 12 IHCs per cochlea, *n* = 4-6. Data shown as means ± SEM, \**P* < 0.05, \*\**P* < 0.01, \*\*\**P* < 0.001 by one-way ANOVA with Bonferroni post hoc test.

### KSR1 KO mice are resistant to ERK activation following cisplatin administration and noise exposure

Previous studies indicate cisplatin administration and noise exposure induces phosphorylation of ERK1/2 in the cochlea^6, 8,10,12^. To determine whether KSR1 mice are resistant to cisplatin and noise activation of ERK1/2, cochleae from carrier alone or cisplatin treated WT and KO mice were collected 45 minutes after cisplatin treatment. Another group of WT and KO mice were exposed to noise and their cochlea were collected immediately following a 100 dB SPL (8-16 kHz) noise exposure for 2 hours. Cross-sections of cochleae were then stained with antibodies for pERK1/2, neuronal marker Tuj1, and DAPI for cell nuclei and imaged by confocal microscopy (Figure 7). Representative images show cisplatin treated and noise exposed WT mice have increased pERK1/2 in the organ of Corti region compared to carrier alone treated mice. Conversely, no difference in pERK1/2 staining was observed between carrier alone, cisplatin treated, and noise exposed KSR1 KO mice. Additionally, while the stria vascularis was also analyzed, no pERK staining was observed in any of the treatments and KSR1 genotypes.

**Figure 7:**
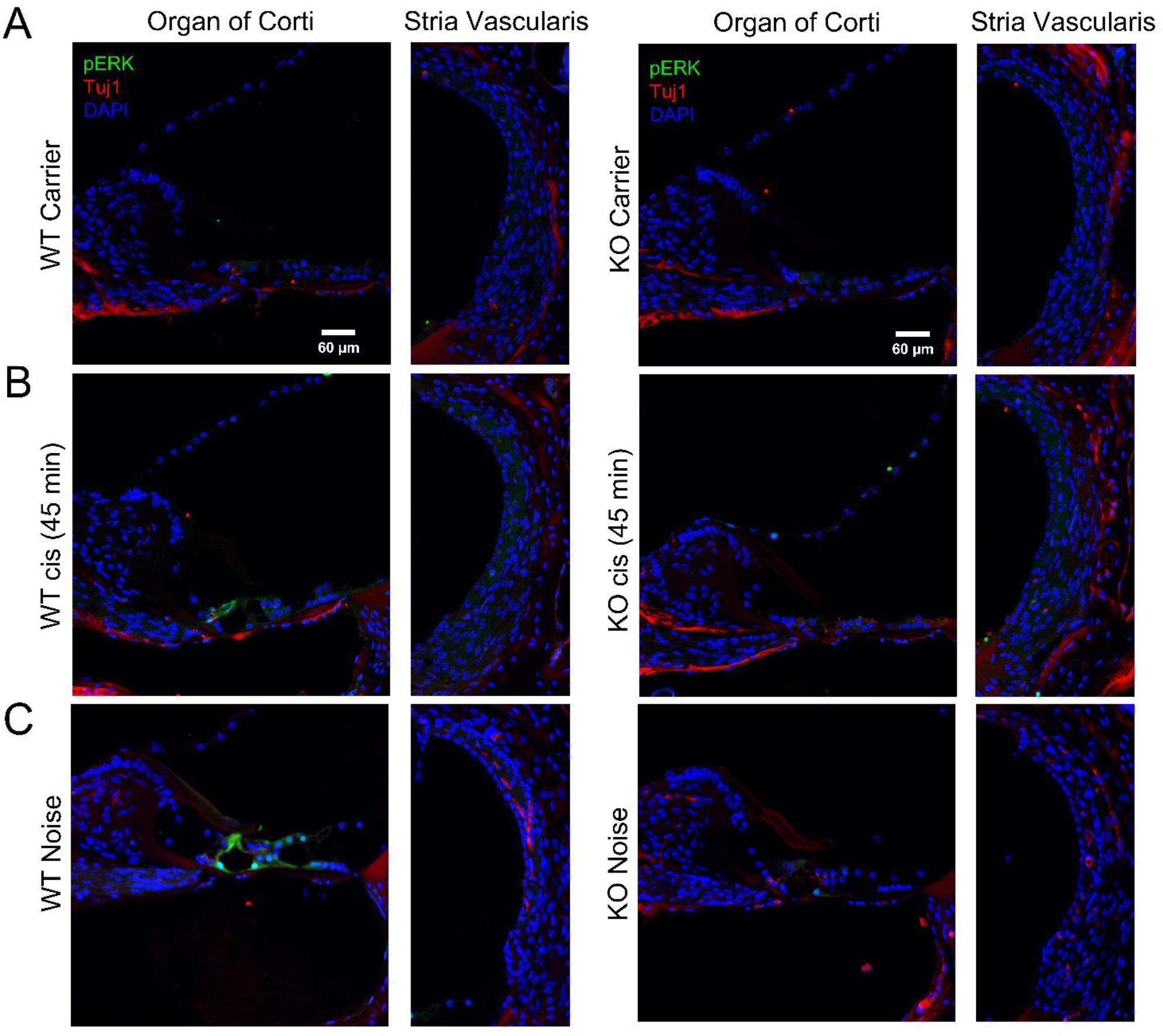
Cisplatin and noise exposure induces ERK phosphorylation in KSR1 WT, but not KO mice. Representative confocal images of the organ of Corti and Stria Vascularis of cochleae collected from KSR1 WT and KO mice 45 minutes after treatment with **(A)** carrier alone or **(B)** 18 mg/kg cisplatin IP injection. **(C)** Noise exposed KSR1 WT and KO cochleae were collected immediately following a 100 dB (8-16 kHz) noise exposure for 2 hours. Tissues were stained for pERK (green), Tuj1 (red), and DAPI (blue). Scale Bar = 60µm *n* = 3.

## Discussion

Our laboratory has previously shown that multiple inhibitors of the MAPK pathway protect from cisplatin and noise-induced hearing loss^6–8^. This study is a genetic proof that the pathway is indeed involved in both types of hearing loss and that dabrafenib is exhibiting its protective effect through inhibition of the MAPK pathway, and not through off target effects (Figure 8). The MAPK kinases, BRAF, MEK, and ERK, are all ubiquitously expressed throughout the inner ear and KSR1 is also expressed in many of the inner ear cell types^27^. KSR1 is the dominate isoform expressed rather than KSR2 which is one of the reasons this genetic mouse model is useful for studying the role of the MAPK pathway in hearing loss. Both isoforms can act as scaffolding proteins in the MAPK pathway, but they can have different physiological functions^28,29^. KSR1 was hypothesized to play more of a role in hearing loss than the KSR2 isoform due to KSR1 having much higher expression in the inner ear and being required for optimal MAPK signaling. KSR1 was predominately expressed in OHCs, inner border cells, inner phalangeal cells, fibrocytes, intermediate cells, inner/outer sulcus cells, root/spindle cells, and Reisner’s membrane. We hypothesize that these cell types are critical in the stress response that leads to hearing loss because KSR1 KO mice were resistant to cisplatin and noise-induced hearing loss. Overall, this study shows that knocking out KSR1, as well as inhibiting BRAF, MEK, and ERK, are all promising therapeutic targets to protect from both cisplatin and noise-induced hearing loss.

**Figure 8:**
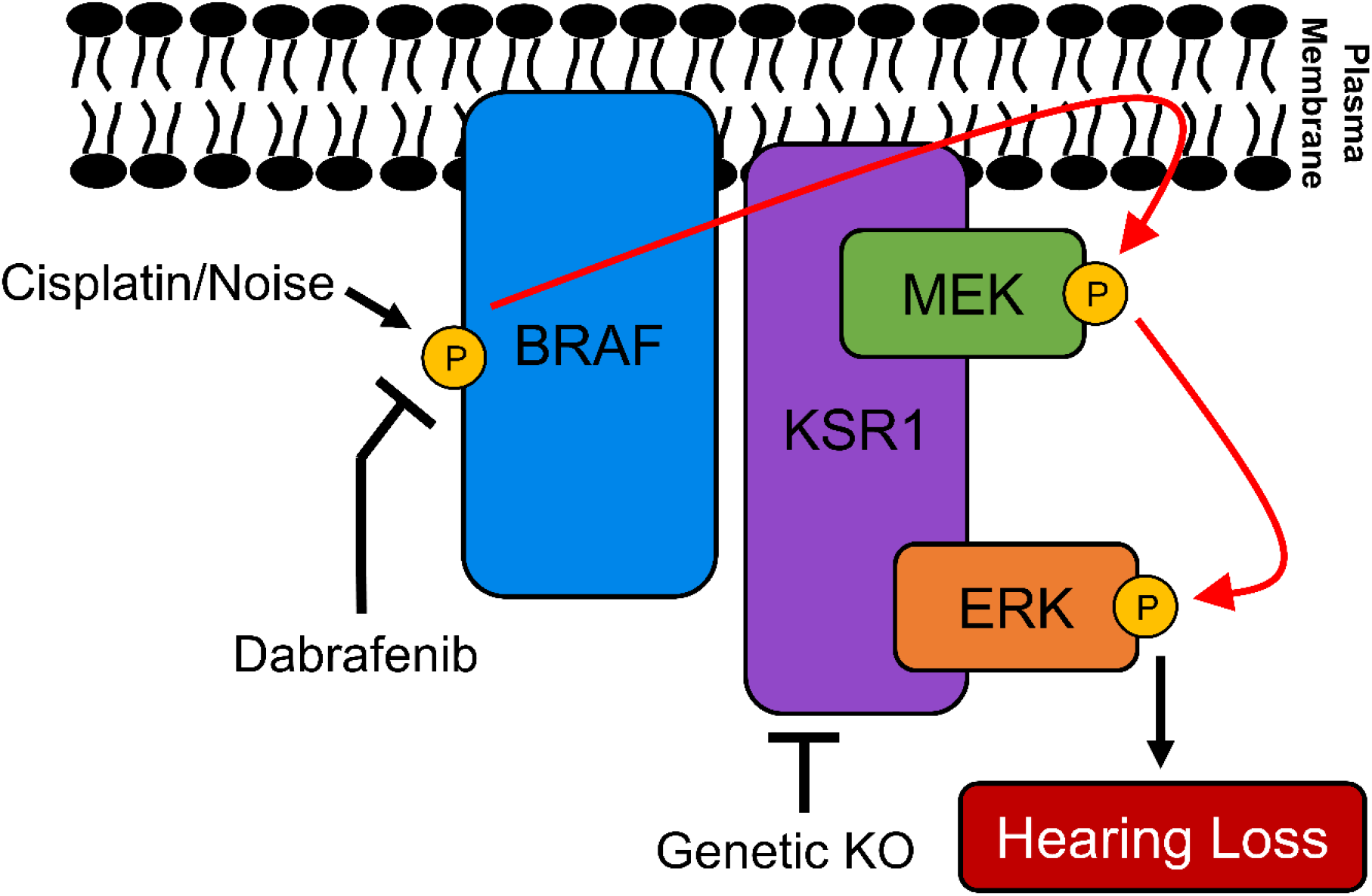
Induction of the putative BRAF/ERK/MEK phosphorylation cascade leads to hearing loss in mice. Cisplatin or noise exposure leads to phosphorylation and activation of BRAF at S445. KSR1, a scaffolding protein which binds MEK and ERK in the cytoplasm, then dimerizes with activated BRAF on the plasma membrane to colocalize all three kinases. BRAF then phosphorylates and activates MEK1/2 at S217/221, and MEK phosphorylates and activates ERK1/2 at T202/Y204 which leads to permanent hearing loss. In the present study, we demonstrate disruption of the MAPK pathway, by inhibition of BRAF with dabrafenib or genetic KO of KSR1, significantly reduces hearing loss caused by cisplatin or noise exposure

KSR1 KO mice were shown to have significantly less hearing loss, as demonstrated by ABR, DPOAE, outer hair cell counts, and ctbp2 staining, compared to their wild-type littermates. KSR1 KO mice have no difference in baseline hearing compared to WT mice which demonstrates that eliminating the protein does not affect normal hearing but is important in the damage process. KO mice were almost completely resistant to cisplatin ototoxicity and offered approximately 80% protection from noise-induced hearing loss. The level of protection following 106 dB SPL noise exposure is approximately 53% of the 100 dB 2-hour noise protection but that is expected due to the higher noise intensity, and this has been demonstrated with pharmacological inhibition of the pathway^7^. Pharmacological inhibition of BRAF, MEK, and ERK all significantly protect from cisplatin and noise-induced hearing loss and this genetic knockdown of the MAPK pathway confers similar protection as the kinase inhibitors previously studied^6–8^.

Previously, our laboratory has demonstrated that dabrafenib, a BRAF inhibitor, significantly protects multiple different strains of mice from cisplatin ototoxicity and noise-induced hearing loss^6,8^. This study confirms that dabrafenib protects from both types of hearing loss through inhibition of the MAPK pathway, and not by off target effects. If dabrafenib was acting through an off-target pathway, KO mice treated with the drug were expected to have different levels of hearing protection compared to KO mice not treated with the drug, which was not observed (Figures 4 & 6). Oral administration of dabrafenib to KSR1 KO mice did not confer any extra protection compared to KO mice not treated with the drug. WT KSR1 mice treated with dabrafenib offered the same amount of protection as both KO KSR1 groups from cisplatin and noise-induced hearing loss; therefore, dabrafenib is protecting through the MAPK pathway. This was observed following both cisplatin administration and noise exposure. It is crucial to first demonstrate the mechanistic target of the drug dabrafenib to initiate and design clinical trials for protection from cisplatin-induced hearing loss^3^.

Along with previously studied FVB and CBA mice, WT C57BL/6 mice had significant increases in ERK phosphorylation following cisplatin and noise exposure which shows that the MAPK pathway is activated following these stimuli; however, KSR1 KO mice were resistant to this increase in pERK (Figure 7)^6,8^. KO mice also had significantly lower amounts of MAPK activation at baseline levels compared to WT littermates as shown by the western blot data in Figure 2. Genetic KO of the KSR1 gene did not completely abolish MAPK activation, but significantly knocked it down which is crucial for the feasibility of our present studies. The MAPK pathway is critical in a multitude of cellular processes and complete knockout of the pathway could be lethal to the inner ear cells^30,31^.

Overactivation of the MAPK pathway, as observed with cisplatin and noise exposure in the inner ear, can lead to downstream cellular stress pathways and eventually to cell death and dysfunction ^6,9–11,23,32^. Stimulation of the MAPK pathway has previously been associated with cell survival and proliferation; however, our studies here and previous studies have suggested that this pathway can also act as a cellular stress pathway in some non-proliferating cells such as the inner ear and the kidney^6,8,24^. Following cisplatin and noise exposure, there is a significant increase in ROS, proinflammatory cytokines and chemokines, and other inflammatory markers, which are all markers of cell stress^33–36^. Inhibition of the MAPK pathway could be lowering the amount of these cell stress molecular events which would lead to less cellular dysfunction and death. Recently, our laboratory has shown that ERK1/2 inhibition following noise exposure lowered the number of infiltrating immune cells^7^. Immune cell infiltration and inflammation has been proposed to exacerbate the hearing loss that occurs following cisplatin administration and noise exposure^33,37–40^. Lowering MAPK activation following these insults could decrease the number of infiltrating immune cells which could lower the amount of hearing loss. These are possible mechanisms through which MAPK inhibitors and KO of the KSR1 gene is protecting from hearing loss, and our future studies will investigate this topic further.

Interestingly, in Figure 7, phosphorylated ERK was increased following cisplatin and noise exposure; however, the pERK staining was more intensive after noise exposure compared to cisplatin administration. Noise exposure in our experiments is an intense two hours, acute damaging stimulus while in cisplatin administration, the damaging stimulus accumulates over time as cisplatin enters and accumulates in the ear^41–43^. This could be the reason why pERK activation is greater in intensity following noise exposure in our experiments compared to cisplatin administration. The time point we chose to test ERK phosphorylation could also be a reason for the difference in pERK signal because cisplatin damage may need to accumulate longer to see stronger signals^41^. Remarkably, activation of the MAPK pathway was shown to occur after both cisplatin administration and noise exposure in the same cells in the cochlea, which shows that there are common similarities in the mechanisms that lead to both types of hearing loss^6,8,44^.

Activation of the MAPK pathway as a stress pathway following various damaging stimuli could be a general cell stress pathway that is activated in different postmitotic cells, not just the inner ear cells. Other studies, including ours, have shown that activation of MAPK proteins in other tissues, such as the kidneys and neurons, leads to cellular stress and death^24,25,32,45,46^. This current study contributes further evidence that the MAPK pathway can act as a stress response pathway and targeting the proteins in the pathway could be beneficial for preventing other types of diseases such as acute kidney injury and neuroinflammatory disorders^23–25,32^.

In summary, this study provides clear evidence that activation of the MAPK pathway leads to cisplatin and noise-induced hearing loss and targeting this pathway is a promising therapeutic approach to prevent both types of hearing loss. KSR1 KO mice were resistant to both cisplatin ototoxicity and noise-induced hearing loss while also being resistant to ERK activation, which was highly activated in WT littermates. The KSR1 KO mouse model can be utilized in the future for further mechanistic studies to determine how inhibition of MAPK proteins is leading to protection from hearing loss and for other types of diseases and disorders in which the pathway is activated. Additionally, dabrafenib was shown to be exhibiting its protective effect through the MAPK pathway and not through other off-targets. The MAPK pathway is a promising therapeutic target for preventing cisplatin and noise-induced hearing loss and dabrafenib is a drug that can be utilized to protect from both forms of acoustic trauma.

## Materials and Methods

### Mouse model

Heterozygous C57BL/6 KSR1 WT/KO breeding mice were provided by Dr. Robert Lewis (University of Nebraska Medical Center, Epply Cancer Institute, Nebraska, USA), bred in the animal facility at Creighton University, and used at 6 weeks of age for experiments. Animals were anesthetized by Avertin (2,2,2- tribromoethanol) via intraperitoneal injection at a dose of 500 mg/kg, and complete anesthetization was determined via toe pinch. For all experiments, homozygous WT and KO mice were randomly assigned to experimental groups, maintaining a balance of males and females in each group.

### Single-Cell RNA Sequence Data Analysis

The scRNA dataset previously published by Xu et. al. was analyzed for the expression of MAPK genes in P28 C57BL/6 mice^27^. Briefly, the Seurat objects were processed with the “Read10×” function, and the gene expression data from individual samples were converted into a natural logarithm and normalized under the same condition. The top 2,000 highly variable genes (HVGs) from the normalized expression matrix were identified for further principal component analysis (PCA). The integrated scRNA-seq data assay was created following the Seurat integration procedure. The clustering analysis was performed based on the individual data (P28). We then visualized the cell clusters on the 2D map produced with the UMAP method (Figure 1A). Clusters were primarily annotated using SingleR for a reference-based scRNA-seq cluster annotation; differentially expressed genes (DEGs) with high discrimination abilities were then identified with the FindAllMarkers function. The cluster annotation correction was performed based on DEGs and the well-known cellular markers for cochlear cells. For sub-clustering analysis of all HCs, similar procedures including variable genes identification, dimension reduction, and clustering identification were applied. The cluster-specific overrepresented Gene Set Enrichment Analysis (GSEA) biological process analysis was performed using the clusterProfiler package (version 4.0.5) based on the DEGs in the specific cell cluster compared to other remaining clusters in each dataset. The dataset was then analyzed for expression of MAPK genes including: Braf, Map2k1, Map2k2, Mapk3, Mapk1, Ksr1, and Ksr2. MAPK gene expression patterns were organized by UMAP and violin plots (Figure 1B-C).

### Single dose cisplatin treatment in mice

10 milligrams of cisplatin (479306, Sigma-Aldrich) powder was dissolved in 10 mL of sterile saline (0.9% NaCl) at 37°C for 40 to 60 minutes. 30 mg/kg was administered once to mice via intraperitoneal injection on day 1 of the protocol (Figure 3A). One day before cisplatin injection, mice received 1 mL of saline by subcutaneous injection and were given 1 mL of saline twice a day throughout the protocol until body weight started to recover. The cages of cisplatin treated mice were placed on heating pads until body weights began to recover. Food pellets dipped in DietGel Boost® were placed on the cage floor of cisplatin-treated mice. DietGel Boost® (72-04-5022 Clear H_2_O) is a high calorie dietary supplement that provides extra calorie support for mice. The investigators and veterinary staff carefully monitored for changes in overall health and activity that may have resulted from cisplatin treatment.

### Noise Exposure in Mice

Mice were placed in individual cages in a custom-made acrylic chamber. The sound stimulus was produced by System RZ6 (TDT) equipment and amplified using a 75-A power amplifier (Crown). Sound was delivered to the acrylic chamber via a speaker horn (JBL). The SPL was calibrated with a 1/4-inch free-field microphone (PCB). Before experimental noise exposure, four quadrants of the cage inside the chamber were sampled with the 1/4-inch microphone to ensure that the SPL varied by <0.5 dB across the measured positions. Adult mice were then exposed to 2 h of octave band noise (8–16 kHz) at 100 or 106 dB.

### Compound administration by oral gavage

The compound dabrafenib mesylate was purchased from MedChemExpress and administered to mice via oral gavage. Dabrafenib was dissolved in a mixture of 10% DMSO, 5% Tween 80, 40% PEG-E-300, and 45% saline. For the single dose cisplatin and noise exposure experiments experiment, 12 mg/kg dabrafenib was given to mice twice daily, once in the morning and once at night. For cisplatin experiments, dabrafenib treatment began 45 minutes prior to cisplatin injection (Figure 5A). For noise exposure experiments, dabrafenib treatment began 24 hr after noise exposure (Figure 7A). Treatment continued for a total of 3 days.

### ABR threshold and wave 1 amplitude measurements

ABR waveforms in anesthetized mice were recorded in a sound booth by using subdermal needles positioned in the skull, below the pinna, and at the base of the tail, and the responses were fed into a low-impedance Medusa digital biological amplifier system (RA4L; TDT; 20-dB gain). At the tested frequencies (8, 16, and 32 kHz), the stimulus intensity was reduced in 10-dB steps from 90 to 10 dB to determine the hearing threshold. ABR waveforms were averaged in response to 500 tone bursts with the recorded signals filtered by a band-pass filter from 300 Hz to 3 kHz. ABR threshold was determined by the presence of at least 3 of the 5 waveform peaks. Baseline ABR recordings before any treatment were performed when mice were 6 weeks old. All beginning threshold values were between 10 and 40 dB at all tested frequencies. Post-experimental recordings were performed 14 days following cisplatin treatment or noise exposure. All thresholds were determined independently by two-three experimenters for each mouse who were blind to the treatment the mice received. ABR wave one amplitudes were measured as the difference between the peak of wave 1 and the noise floor of the ABR trace.

### DPOAE measurements

Distortion product otoacoustic emissions were recorded in a sound booth while mice were anesthetized. DPOAE measurements were recorded using the TDT RZ6 processor and BioSigTZ software. The ER10B+ microphone system was inserted into the ear canal in way that allowed for the path to the tympanic membrane to be unobstructed. DPOAE measurements occurred at 8, 16, and 32 kHz with an f2/f1 ratio of 1.2. Tone 1 was *.909 of the center frequency and tone 2 was *1.09 of the center frequency. DPOAE data was recorded every 20.97 milliseconds and average 513 times at each intensity level and frequency. At each tested frequency, the stimulus intensity was reduced in 10 dB steps starting at 90 dB and ending at 10 dB. DPOAE threshold was determined by the presence an emission above the noise floor. Baseline DPOAE recordings occurred when mice were 6 weeks old and tested again 14 days after noise exposure. DPOAE threshold shifts were determined by subtracting the baseline DPOAE recording from the post experimental recording.

### Tissue preparation, immunofluorescence, and Ctbp2/OHC counts

Cochleae from adult mice were prepared and examined as described previously^6,8,44^. Cochleae samples were stained with Hematoxylin and Eosin (H&E) or immunostained with DAPI (1:1000; D1306, Invitrogen), anti-myosin VI (1:400; 25-6791, Proteus Bioscience), anti-Ctbp2 (1:400; 611576, BD Transduction Laboratories), pERK antibody (1:400; 9101L, Cell Signaling), or anti-Tuj1 (1:400; 801201, BioLegend) with secondary antibodies purchased from Invitrogen coupled to anti-rabbit Alexa Fluor 488 (1:400; A11034) or anti- mouse Alexa Fluor 647 (1:800; A32728). All images were acquired with a confocal microscope (LSM 700 or 710, Zeiss). Outer hair cell counts were determined by the total amount of outer hair cells in a 160 µm region. Counts were determined for the 8, 16, and 32 kHz regions. For Ctbp2 puncta counts, the average number of puncta from 12 IHCs in the 16 kHz region was recorded. Cochleae from each experimental group were randomly selected to be imaged for outer hair cell and Ctbp2 counts.

### Immunoblot

Cochleae were rapidly dissected on ice to isolate the organ of Corti, which was homogenized to prepare lysates in lysis buffer (9803; Cell Signaling Technology) after adding protease and phosphatase inhibitors (Roche). Each sample consisted of the combined organ of Corti lysates of both cochleae per mouse. The lysates were centrifuged for 15 min at 15,000 *g* at 4°C, and the supernatants were collected. Protein concentrations were determined with the BCA protein assay kit (23235; Thermo Scientific). 40 µg of protein lysate was loaded on 4–20% SDS-PAGE gels. The following antibodies were used for immunoblot analysis: anti-BRAF (14814S), anti-p-BRAF (Ser445, 2696S), anti-ERK1/2 (4695), anti-p-ERK1/2 (Thr202/Tyr204, 9101S), anti-MEK1/2 (9122S), and anti-p-MEK1/2 (Ser217/221, 41G9S) were obtained from Cell Signaling Technologies, and anti-α Tubulin (T9026) from Millipore sigma. The antibodies were used at dilutions ranging from 1:250 to 1:1,000. Blot intensities were determined using NIH ImageJ software. For KSR1 immunoblot, the organ of Corti from three WT/WT and three KO/KO mice were combined to prepare one lysate sample for each genotype. 100 µg of protein lysate was loaded on 4–20% SDS-PAGE gels. The following antibodies were used for immunoblot analysis: anit-KSR1 (611576) from BD Transduction Laboratories and anti–β-actin (C4; SC- 47778) from Santa Cruz. The antibodies were used at dilutions ranging from 1:250 to 1:1,000.

### Statistical Analysis

Statistics were performed using Prism (GraphPad Software). One-way or two-way analysis of variance (ANOVA) with Bonferroni post hoc test was used to determine mean difference and statistical significance. The student’s T-test was performed when only two values were compared.

### Study Approval

All animal experiments included in this study were approved by Creighton University’s Institutional Animal Care and Use Committee (IACUC) in accordance with policies established by the Animal Welfare Act (AWA) and Public Health Service (PHS).

## Author Contributions

T.T. conceived the project and acquired the funding. M.A.I. performed the cisplatin experiments. M.A.I., R.D.L., R.G.K., and D.F.K. performed the noise exposure experiments. M.A.I. performed the cochlear dissections, outer hair cell, and Ctbp2 counts. M.A.I., R.D.L., D.F.K., and J.M. analyzed ABR and DPOAE data. M.A.I. and R.G.K. performed and imaged the cochlear cryosections. M.A.I. performed and analyzed the immunoblots. J.Z., R.Q., and M.A.I. analyzed the single cell RNA sequencing dataset. M.A.I., R.D.L., and T.T. wrote the manuscript with input from all authors.

## Data, Materials, and Software Availability

All data needed to evaluate the conclusions in the paper are present in the paper. Additional data related to this paper may be requested from the authors.

## Acknowledgements

We thank Kristina Ly, Dr. Christy Howe, Dr. Janee Gelineau-van Waes, Pat Steele, Ann Bryen and the Creighton University ARF staff for assistance with the mouse studies. We thank Emily Schmidt for assisting with collecting data. The research was funded by the National Institutes of Health NIDCD grant 1R01DC018850, American Hearing Research Foundation 2020 grant to T. Teitz, and NIH 1F32DC020102 grant to M.A. Ingersoll. This investigation was conducted in facilities constructed with support from Research Facilities Improvement Program (G20 RR024001-01) from the National Center for Research Resources, NIH. The research was partially conducted at the Auditory and Vestibular Technology Core (AVT) at Creighton University, Omaha, NE (RRID:SCR_023866). This facility is supported by the Creighton University School of Medicine and grants GM103427 and GM139762 from the National Institute of General Medical Science (NIGMS), a component of the National Institutes of Health (NIH). IBIF was constructed with support from grants from the National Center for Research Resources (RR016469) and the NIGMS (GM103427). This investigation is solely the responsibility of the authors and does not necessarily represent the official views of the National Center for Research resources, NIGMS or NIH.

